# Salamander retinal ganglion cell responses to rich stimuli

**DOI:** 10.1101/2020.01.14.906149

**Authors:** Kolia Sadeghi, Michael J. Berry

**Affiliations:** Program in Applied and Computational Mathematics, Princeton University, Princeton, New Jersey, 08544; Department of Molecular Biology, Princeton University Washington Rd., Princeton, New Jersey, 08544-1014

## Abstract

The retina’s phenomenological function is often considered to be well-understood: individual retinal ganglion cells are sensitive to a projection of the light stimulus movie onto a classical center-surround linear filter. Recent models elaborating on this basic framework by adding a second linear filter or spike histories, have been quite successful at predicting ganglion cell spikes for spatially uniform random stimuli, and for random stimuli varying spatially with low resolution. Fitting models for stimuli with more finely grained spatial variations becomes difficult because of the very high dimensionality of such stimuli. We present a method of reducing the dimensionality of a fine one dimensional random stimulus by using wavelets, allowing for several clean predictive linear filters to be found for each cell. For salamander retinal ganglion cells, we find in addition to the spike triggered average, 3 identifiable types of linear filters which modulate the firing of most cells. While some cells can be modeled fairly accurately, many cells are poorly captured, even with as many as 4 filters. The new linear filters we find shed some light on the nonlinearities in the retina’s integration of temporal and fine spatial information.

## Introduction

Much fruitful effort has been devoted to modeling the various stages of the signaling cascade in the retina, focusing on photoreceptors, bipolar cells, ganglion cells, and other cell types, uncovering several adaptive and nonlinear effects. On the other hand, phenomenological studies aiming at predicting the responses of retinal ganglion cells to general stimuli directly, that is without explicitly modeling the various stages of the signaling cascade, have only strayed a little from the basic classical view of a receptive field organized as a center and a surround, wherein spatial information is summed mostly linearly. A notable exception to this are studies introducing rectifying spatial subunits (Hochstein & Shapley, 1976; Shapley & Victor, 1979; Ölveczky, Baccus, & Meister, 2003; Baccus, Ölveczky, Manu, & Meister, 2008), although the stimuli used were sinusoidal waves and drifting gratings, which are not spatially rich. For tractability, most recent phenomenological approaches aiming at predicting responses to rich stimuli have elaborated on the simple linear-nonlinear (LN) model class, where spiking probabilities are some nonlinear function of a projection onto one or two linear filters of the stimulus and possibly past spikes; for reviews, see (Paninski, 2004; Simoncelli, Paninski, Pillow, & Schwartz, 2004; Truccolo, Eden, Fellows, & Donoghue, 2005). This approach has been quite successful at predicting ganglion cell spiking probabilities for spatially uniform random stimuli, by using covariance analysis to find more than one linear filter (Bialek & Steveninck, 2005; Fairhall et al., 2006). Belonging to the same LN model class, Generalized Linear Models (GLM), with and without taking into account spiking histories of a population of cells, have been shown to be excellent predictors of macaque parasol cell responses to spatio-temporal random stimuli, albeit when spatial pixels have sizes comparable to receptive field center widths (Pillow et al., 2008).

Nonetheless, the ultimate goal of modeling retinal responses to general random stimuli has remained elusive for stimuli with spatial variations on a scale much finer than receptive field centers, and for all ganglion cell types; this is because typical experimental data is insufficiently long to identify and fit models.

We present a method of reducing the dimensionality of a fine flickering strips stimulus by using wavelets. We then proceed to fit various models using wavelet coefficients to represent past stimuli and compare the amount of information they capture to a tight upper information bound. First, it is noted that models based only on the spike triggered average (STA) do very poorly at predicting responses for many cells. We augment the STA models with additional linear filters found using covariance analysis. Even the augmented models fall short of fully capturing ganglion cell light responses for every cell. Nonetheless, the new filters found by covariance analysis give some insight into visual features other than the classical receptive field which influence ganglion cell responses. Models incorporating past spike histories into covariance based and GLM models are shown in supplements to give informative but very redundant filters.

## Materials and Methods

### Electrophysiological Recording

Retinae from larval tiger salamanders (*Ambystoma tigrinum*) were isolated from the eye retaining the pigment epithelium, placed over a multi-electrode array and perfused with room temperature Ringers oxygenated medium. Extracellular voltages recorded by a MultiChannel Systems MEA 60 microelectrode array were streamed to disk for off-line analysis at 10 kSamples/s/channel (Segev, Goodhouse, Puchalla, & Berry, 2004). Spike waveforms were sorted using the spike size and shape from the full waveform on 30 electrodes.

### Visual Stimulation and cross-validation

Frames consisting of 200 black or white 22 *µ*m wide vertical strips were pseudorandomly reconstructed and projected every 16 *ms* from a computer monitor onto the photoreceptor layer. Each strip had a 50% chance of being black or white, independently of all other strips. Two separate experiments were run. The first experiment (37 cells) consisted in 43 minutes of random stimulus. For the second experiment (23 cells), the stimulus protocol consisted in 6 runs each consisting in 20 repeats of the same 30 second stretch of random stimulus, followed by 20 minutes of novel random stimulus. For the second experiment, only the last 4 runs were kept for analysis, as the firing rates of many cells changed during the first two runs.

For model fitting and evaluation, each of our two data sets was partitioned into a training set, a validation set, and a testing set. The testing set was set aside for evaluation purposes only: all estimates of mutual information (in *bits/spike*) captured by our various models for the first (second) experiment were calculated using 16 (8) minutes of recordings otherwise unused for any other purpose. Training sets 16 (8) minutes long were used for finding STAs, covariance analysis, fitting GLM model parameters, and estimating nonlinearities. Validation sets 8 (4) minutes long were used to calculate gaussian scores in order to select informative covariance filters, and to find optimal regularization parameters for our GLM models.

### Past stimuli and spike trains

Our models predict the firing probability of a given cell at time *t* as a function of the past stimulus and (optionally) past spike trains. The stimulus leading up to time *t* was characterized by the spatial light intensity at 128 time points equally spaced between the spike time and 512 *ms* before the spike, corresponding to a temporal resolution of 4 *ms*. For each cell, the spatial domain considered was restricted to 128 strips. Thus the stimulus leading to time *t* was restricted to be a 2-D vector of 128*128 binary light intensities -either black or white - with one dimension of the array representing time sampling, and the other representing strips. Some of our models also depend on the past spikes of the cells in the population (see and); for these, the spike train leading up to time *t* was characterized by the vector of times since the last spike of each cell.

### Wavelet analysis of stimulus pasts

The receptive fields of retinal ganglion cells tend to be localized in space and time, and they tend to be smooth. An efficient tool for analyzing and representing data with such properties is well known: wavelets. Using a wavelet decomposition of the stimulus, we kept only those wavelet coefficients which were deemed informative about a cell’s activity. Using wavelet software detailed in (Selesnick, 2004), a 5-fold redundant wavelet basis was used to decompose the 128*128 array of binary light intensities leading to any time of interest *t*. A wavelet coefficient was deemed significantly informative about a cell’s spike times when its mean and mean square value were significantly different at spike times than at uniformly sampled times.

More precisely for a fixed wavelet, let’s write 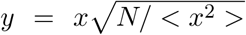, where *x* is the wavelet’s coefficient, < *x*^2^ > denotes the mean squared coefficients for the wavelet at uniformly random times, and <. >_*spike*_ denote averages over *N* spike times. Then for a perfectly uninformative wavelet:

- < *y* >_*spike*_ has an approximately zero mean unit variance normal distribution.
- < *y*^2^ >_*spike*_ has an approximately *χ*^2^ distribution with *N* degrees of freedom.
- 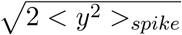 is approximately normally distributed with mean 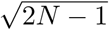 and unit variance, and is independent from < *y* >_*spike*_.
- 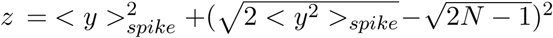 is approximately *χ*^2^ with 2 degrees of freedom.

A wavelet was deemed significant if *z* was larger than the inverse cumulative distribution function (*icdf*) of the *χ*^2^(2) distribution at probability 1 − 0.01*/N*_*wavelets*_:

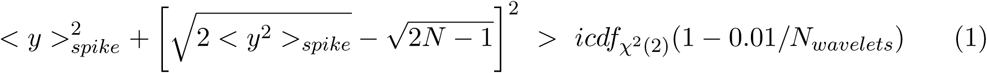

In other words wavelets were deemed significant if renormalized gaussianized versions of their mean and mean square statistics jointly deviated significantly from chance at spike times (p-value < 0.01*/N*_*wavelets*_ where *N*_*wavelets*_ is the total number of wavelets, Fig. 1). This significance level was chosen to be stringent enough for irrelevant wavelets to be discarded, while being inclusive enough to keep the more informative wavelets. For the first experiment, this resulted in 423-1567 informative coefficients (depending on cells, *n* =37). For the second experiment, wavelets were kept only if 3 out of the 4 periods of 20 minutes of recording were deemed significantly informative, resulting in 281-927 informative coefficients (depending on cells, *n* =23). Note that since the wavelet transform we used was 5-fold redundant, the effective dimensionality of the subspace obtained by this projection is effectively approximately 5 times smaller than the actual number of chosen wavelets. All subsequent stimulus-dependent analysis was done using only this subset of chosen wavelet coefficients. Choosing an overcomplete wavelet basis and a hard cutoff allowed us to avoid hard cutoff artifacts that a non-redundant wavelet basis would have produced, making for a redundant yet informative and manageable representation of past stimuli.

**Figure 1.**
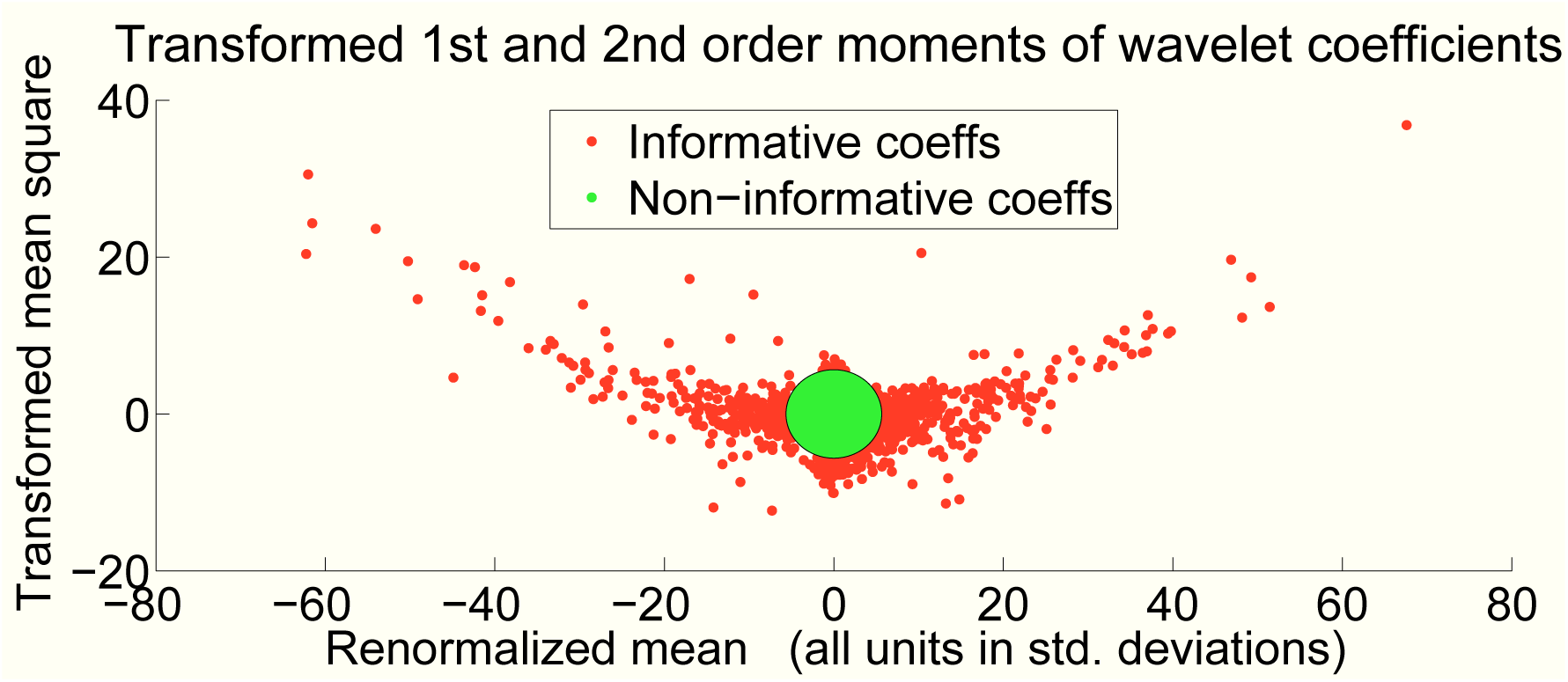
Scatter plot of wavelet coefficients represented by their mean and transformed mean square across spike times. Both axes were renormalized so that both quantities would have a unit standard deviation if averaged over random times. The approximately chi-squared distributed mean squares were transformed to obtain a more gaussian-like null-hypothesis distribution. Wavelet coefficients deviating less than approximately 5 standard deviations from zero are deemed uninformative (green points) and thrown away.

### A common framework: linear-nonlinear models

We predicted ganglion cell spikes by fitting several models. All models were part of the linear-nonlinear family. In such models, a first stage computes *D* linear combinations of the wavelet coefficients characterizing the past (Fig. 2). This is equivalent to convolving the stimulus with *D* linear filters. This linear projection results for each time *t* in a *D*-dimensional vector 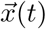 characterizing the past. A second stage produces a probability of spiking by applying a static nonlinearity which collapses 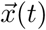 onto a single number *r*(*t*), the predicted spiking intensity. To appropriately model the firing intensity of a cell, the output *r*(*t*) of the nonlinearity must approximate the ideal quantity:

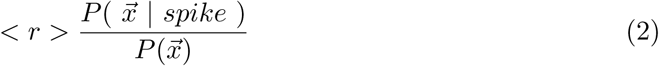

where < *r* > is the mean firing rate of the cell, 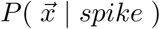 is the probability distribution of 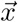 at spike times, and 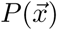 is the probability density of 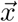 at uniformly random times. This can be seen as a simple application of Bayes’ rule to 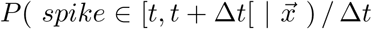 for infinitesimal Δ*t*, noting that < *r* > is the limit when Δ*t* goes to zero of *P* (*spike* ∈ [*t, t* + Δ*t*[) */* Δ*t*.

**Figure 2.**
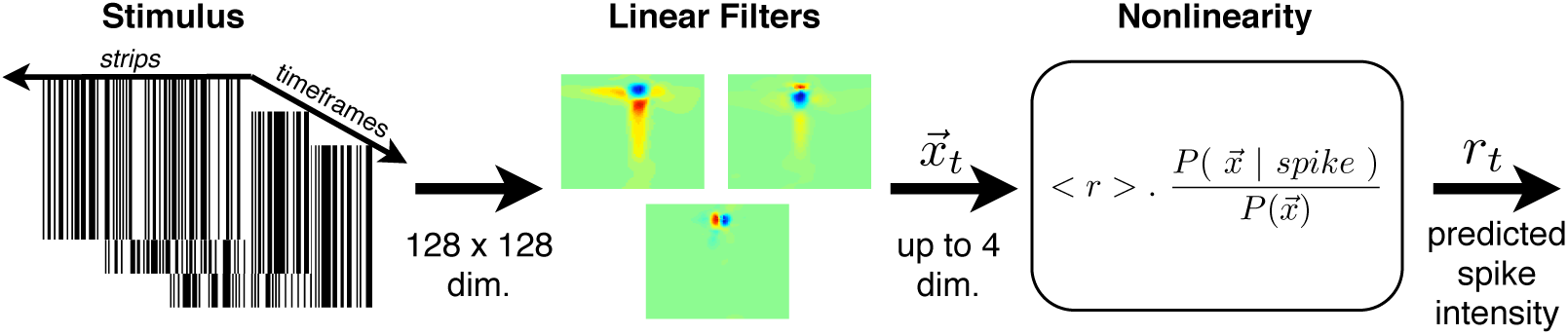
Schematic representation of Linear-Nonlinear models. One or more linear filters are applied to the past stimulus at a given time *t*. The resulting one to four dimensional linear projection 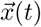 is mapped onto a predicted instantaneous spike rate *r*(*t*) by a function proportional to 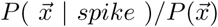, where 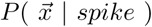 is the probability density of 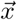 at spike times, and 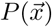 is the probability density of 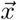 at uniformly random times.

All of the models we tried belong to this linear-nonlinear framework. The number of linear filters used ranged from one to 4. Some models predicted a cell’s activity based on the past spikes of the population of cells and the past stimulus, while others depended on the past stimulus only. Different methods were used to fit the linear filters: Spike Triggered Average, covariance analysis, or Generalized Linear Model fitting. Also, two different methods were used to fit the nonlinearity: direct or factorized gaussian kernel smoothing. All of these different techniques are detailed in section below.

### Covariance analysis

In order to find linear filters which capture information missed by the STA, we performed a covariance analysis on the residual pasts obtained by projecting out the contribution along the STA: each vector 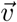 characterizing the past was substituted with 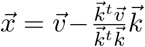, where 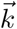 is the STA linear filter. Covariance analysis then allowed us to find additional informative linear filters, in a way resembling (Brenner, Bialek, & Steveninck, 2000; Fairhall et al., 2006). This was done as follows: a set of *N* residual vectors 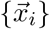 characterizing past stimuli at spike times *i*, and a set of *M* residual vectors 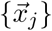 characterizing past stimuli at uniformly random times *j*, were used to calculate the empirical covariance matrices **C**_*spike*_ and **C**_*random*_. These matrices were normalized by dividing by the sum of the variance at spike times and at random times, separately for each dimension. Since the STA direction was projected out, the residuals at spike times and at random times both have zero mean. The covariance analysis then consisted in diagonalizing **C**_*ratio*_, where:

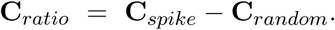

Eigenvectors associated with large positive or negative eigenvalues of **C**_*ratio*_ correspond to directions in which the variance was different at spike times compared to random times, indicating directions that are good candidates for being informative (Steveninck & Bialek, 1988; Fairhall et al., 2006).

The set of eigenvector filters was then ranked for information capture assuming that projections along the filter have gaussian distributions: a gaussian was fit to the projections along each filter at spike times, and another gaussian fit to projections at random times. The average over spike times of the log-ratio of the two densities, the ‘gaussian score’, is a quick approximation of the information captured by each filter. Since both gaussian distributions have zero mean, the gaussian score is written:

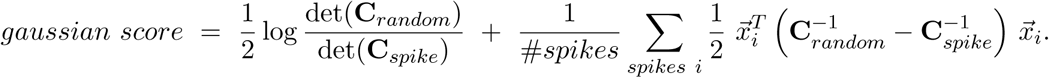

This approximation is justified since in practice the projections of pasts at spike times and random times are indeed close to being gaussian. Eigenvectors were ranked in decreasing order of gaussian score, and the three best filters were kept, provided they had a gaussian score of at least 0.02 bits. All gaussian scores were evaluated using validation data distinct from the training and test data, set aside for this purpose, so as to minimize overfitting.

### Fitting nonlinearities

Once linear filters were found for a cell, we proceeded to fit a nonlinearity which outputs a predicted instantaneous firing rate as a function of the projections 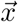 along the filters. As noted above, such a nonlinearity must approximate the instantaneous spiking rate 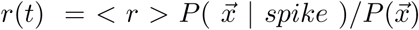. We fit this quantity in a general nonparametric fashion by separately fitting the probability distributions 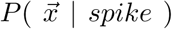 and 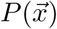 using standard kernel density estimation (KDE) by convolving the cloud of points 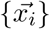 at spike times *j* and the cloud of points 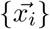 at uniformly random times *i* with a gaussian kernel (Silverman, 1986; Gray & Moore, 2003).

### Measuring informativeness

In order to evaluate models, it is useful to measure how informative a given function 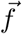 of the past is about a particular cell’s spiking. The quantities 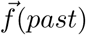 of interest to us are the outputs of one or more linear filters, or of model nonlinearities. A theoretically sound measure of informativeness is the mutual information 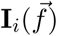, in *bits/spike* between the quantity 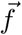 and the spikes of cell *i*. Note that we could have used the likelihood of the spike train given the model instead, as is often used in similar studies. The likelihood is expressible using the spiking intensity *r*(*t*) (often called the conditional intensity) of the model by 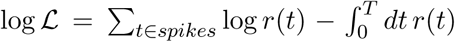. However, the log-likelihood does not scale linearly when time units are changed, and thus cannot be interpreted as measuring bits of information. The mutual information we use does not suffer from this problem, and corresponds to an actual number of bits of information. Furthermore, it allows us to evaluate the information content of linear filters without explicitly fitting a nonlinearity. This mutual information in *bits/spike* is obtained by calculating the Kullback-Leibler divergence between the distributions of 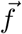 at two ensembles of times: observed spike times, and uniformly random times (Brenner, Strong, Koberle, Bialek, & Steveninck, 2000):

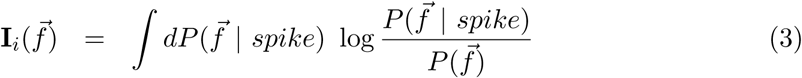

where as before, 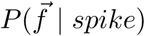 is the probability distribution of 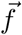 at spike times, while 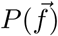 is its probability distribution at uniformly random times. See section for a derivation. Many estimators of Kullback-Leibler divergences have been proposed and studied in the literature. We choose to use a well known estimator based on nearest-neighbor distances (Kozachenko & Leonenko, 1987; Victor, 2002; Perez-Cruz, 2008, 2007). For the quantity of data available, this estimator provides reliable estimates of information for up to four filters. The estimator acts on the collection of values 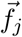 of 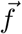 sampled at spike times *j* representing 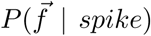 and a collection of values 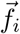 sampled uniformly in time representing 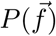. The estimated Kullback-Leibler divergence is taken to be:

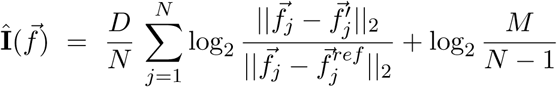

where 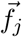 is 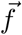 at the *j*-th spike time, 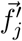 is the sample at another spike time for which the value of 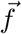 is closest to 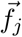, and similarly 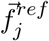 is the closest value to 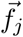 at a reference time sample. Here, ‖·‖_2_ is the usual euclidean distance, *D* is the dimensionality of the data considered, *N* is the total number of spike time samples, and *M* is the total number of reference time samples.

Note that 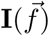 is the mutual information between the stimulus and 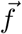 with spike times considered independently, ignoring statistical structure within the spike train itself ; it is not to be confused with the mutual information between the function of time 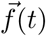 and the whole spike train. We will be able to look at the influence of statistical structure within the spike train itself by including past spikes in our models, i.e. by considering quantities 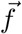 which depend on past spikes as well as the past stimulus (see sections and).

In the case of models which depend only on past stimuli, the information **I** can be compared to a true and easily measurable benchmark which we will call the information bound ℬ. All functions of the past stimulus 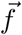, including our model filters and nonlinear output, satisfy 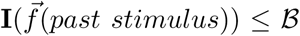, with equality achievable only if 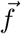 preserves all the information in the past stimulus about individual spikes. The information bound is a measure of the reproducibility of a cell’s responses to a stimulus repeated many times, ignoring interactions between spikes:

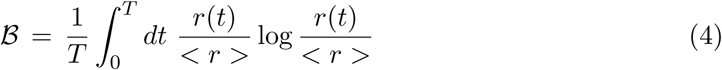

where *T* is the length of the repeated stimulus, *r*(*t*) is the measured firing rate of the cell at time *t* of the repeated stimulus, and < *r* > is the mean firing rate of the cell. We estimated the information bound without binning or otherwise estimating *r*(*t*) by noticing that the quantity *r*(*t*) */* < *r* > can be interpreted as the ratio between the probability densities *r*(*t*) */*(*T* < *r* >) and 1*/T*. The information bound ℬ is then estimated by interpreting it as a Kullback-Leibler divergence and once again using the nearest-neighbor based estimator. The estimator was used directly on spike times elicited by the repeated stimulus as the set of target samples {*t*_*j*_} and uniformly random time points as the reference samples {*s*_*j*_ }, with:

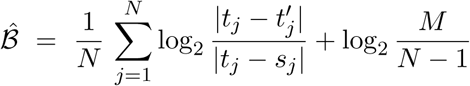

where *t*_*j*_ is the *j*-th spike time, 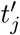 is the (next) closest spike time to *t*_*j*_, and *s*_*j*_ is the closest reference time sample to *t*_*j*_. Here again, *N* is the total number of spike time samples, and *M* is the total number of reference time samples. This estimator provides reliable estimates provided we use large enough *M* so that the density of reference samples is at least as large as the highest local density of samples at spike times.

## Results

### Spike Triggered Stimulus Averages

To illustrate the wavelet analysis performed in stimulus space, we first look at the spike-triggered stimulus average (STA). The STA formed by averaging the raw 128-by-128 binary stimuli before each spike is very noisy, even for cells with thousands of spikes, while its projection onto the wavelets chosen for each cell is much more smooth. As one can see by subtracting the wavelet STA from the raw STA, the wavelet coefficients capture most of the structure of the receptive field, while essentially leaving out noise (Fig. 3).

**Figure 3.**
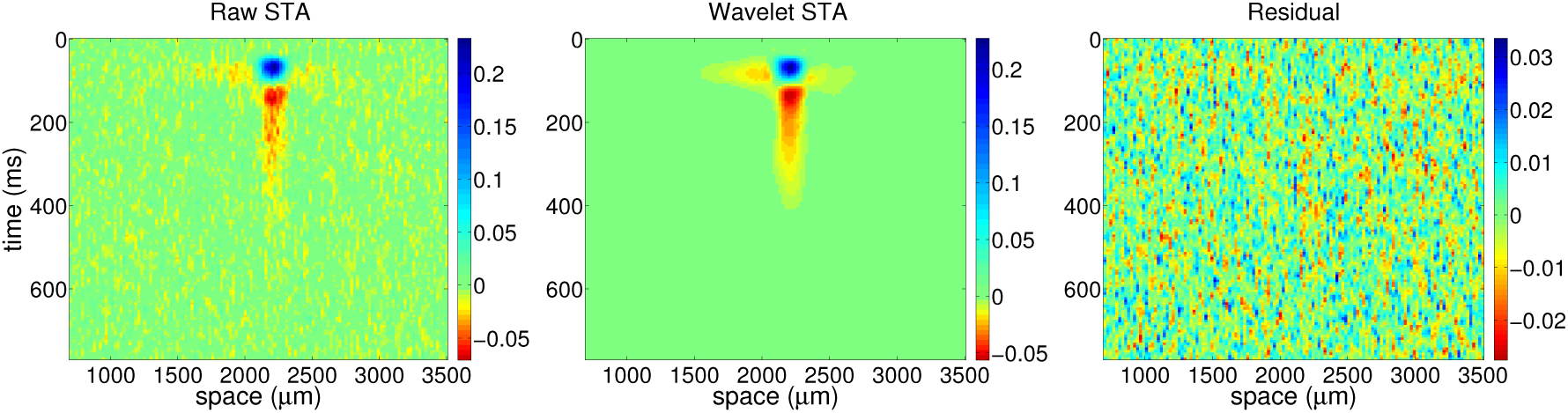
*Left:* Raw Spike Triggered Average. *Middle:* projection onto informative wavelets. *Right:* the residual obtained by subtracting the wavelet projection from the raw STA. Very little low-frequency structure has been missed by the wavelet projection.

The information captured by the STA projected onto the chosen wavelet subspace was significantly larger than the information captured by the raw STA, even after adjusting the raw STA’s spatial extent in pixels to an approximately optimal size by hand. This is simply because the raw STA over-fits the data which it is an average of: as with the rest of this paper, all information estimates cited are obtained from test data not seen during model fitting, and any over-fitting of model parameters result in less information captured on the test data set. Hereafter, we will refer to the STA onto the chosen wavelet subspace simply as ‘the STA’.

Perhaps surprisingly, the STA fares poorly as a predictor of spiking activity (Fig. 4), even once it has been denoised by projecting it onto our wavelet subspace. The proportion of the information bound ℬ captured by the STA was at most 68% (the minimum was 7%), and the proportion of the information bound captured by the STA was less than 32% for more than half of cells. This prompted us to look for additionally informative filters to supplement the STA.

**Figure 4.**
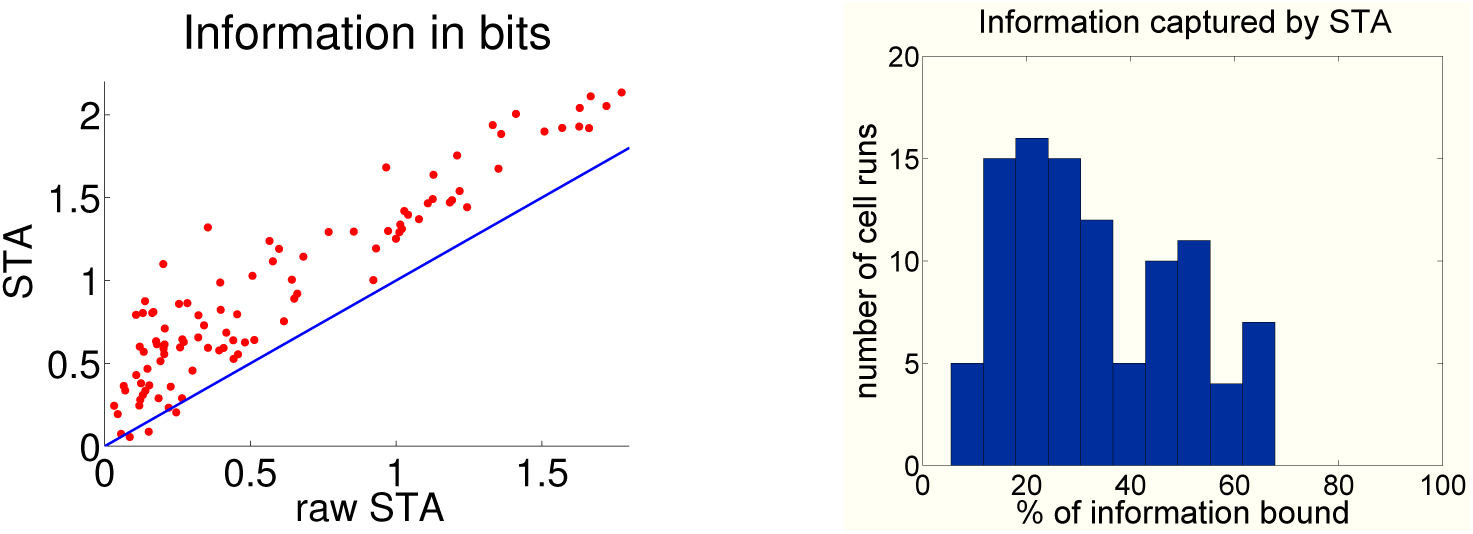
*Left:* Scatter plot of information captured by the STA projected onto chosen wavelets versus onto the raw STA, *n* = 23 cells, for 4 different stimulus segments each. To be fair, since the 128-by-128 area used for our wavelet methods is very large, the raw STA was taken on a smaller area of stimulus space: 50 spatial pixels, and 24 temporal samples spaced by 10 *ms*. This window size was chosen by trial and error to be approximately maximally informative, by enclosing enough of the receptive field without being too large for overfitting to become dominant. *Right:* Histogram of the percentage of information bound ℬ captured by the STA projected onto our chosen wavelets.

### Cell types

For ease of reference in subsequent sections, let us review the range of cell types which can be observed among salamander retina ganglion cells. As in (Segev, Puchalla, & Berry, 2006), the cells we recorded from can be classified into a half-dozen types by looking at the timecourse of the STA at the spatial location where the STA becomes largest in absolute value (Fig. 5). For our purposes here, it will be sufficient to characterize cells as ranging from being brisk to being sluggish. Brisk cells have the fastest rising timecourses, with biphasic-OFF being the most brisk, followed by monophasic-OFF cells. Sluggish cells have slower timecourses, and correspond to all the other cell types. As we will see, brisk cells will be easier to model using our methods than sluggish cells.

**Figure 5.**
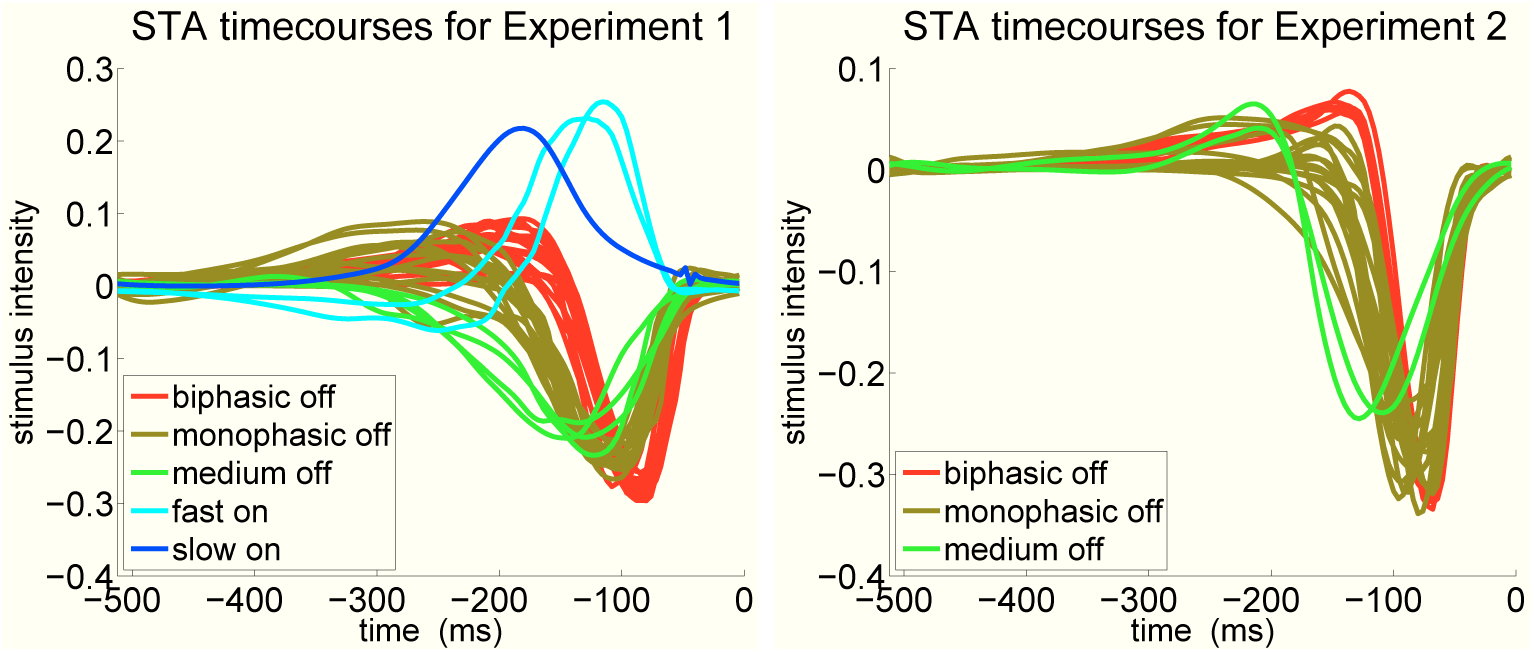
Timecourses of STAs at the location in space where they become maximal. These timecourses can be used to identify ganglion cell types.

### Covariance Analysis features

To further investigate what STA-based models are missing, we used versatile tools which allowed us to find several additional linear filters. We sought linear filters which would supplement the STA filter by first projecting out its contribution ; a covariance analysis was then performed on the residual vectors (see), resulting in up to 3 additional filters for each cell, for a total *D* of up to 4 filters including the STA.

Three new easily recognizable types of spatio-temporal filters emerged from the covariance analysis as being typical features. Unsurprisingly, the most informative feature was still most often, although not always, the STA filter. The three new kinds of filters observed were temporal edges, fast spatial edges and slow spatial edges.

Temporal edge filters (filter 1 in Figures 6 and 7) consist in a lobe close to the time when the STA is maximal, followed in time by a slower lobe of opposite polarity, with both lobes located spatially over the receptive field center: temporal edges are sensitive to temporal changes in light level. In some cases, the slow lobe is flanked by surround effects. Temporal edges are suppressive filters (Rust, Schwartz, Movshon, & Simoncelli, 2004): they have distributions of projections at spike times which have smaller variance than their projections at random times; cells are more likely to fire when the stimulus projection along the temporal edge filter is small. In general, the two lobes of the temporal edge have different sizes: the lobe closest in time to the spike is tighter both in space and in time than the lobe farthest from the spike. Temporal edge filters are vaguely similar to and more informative than the time derivative of the STA.

**Figure 6.**
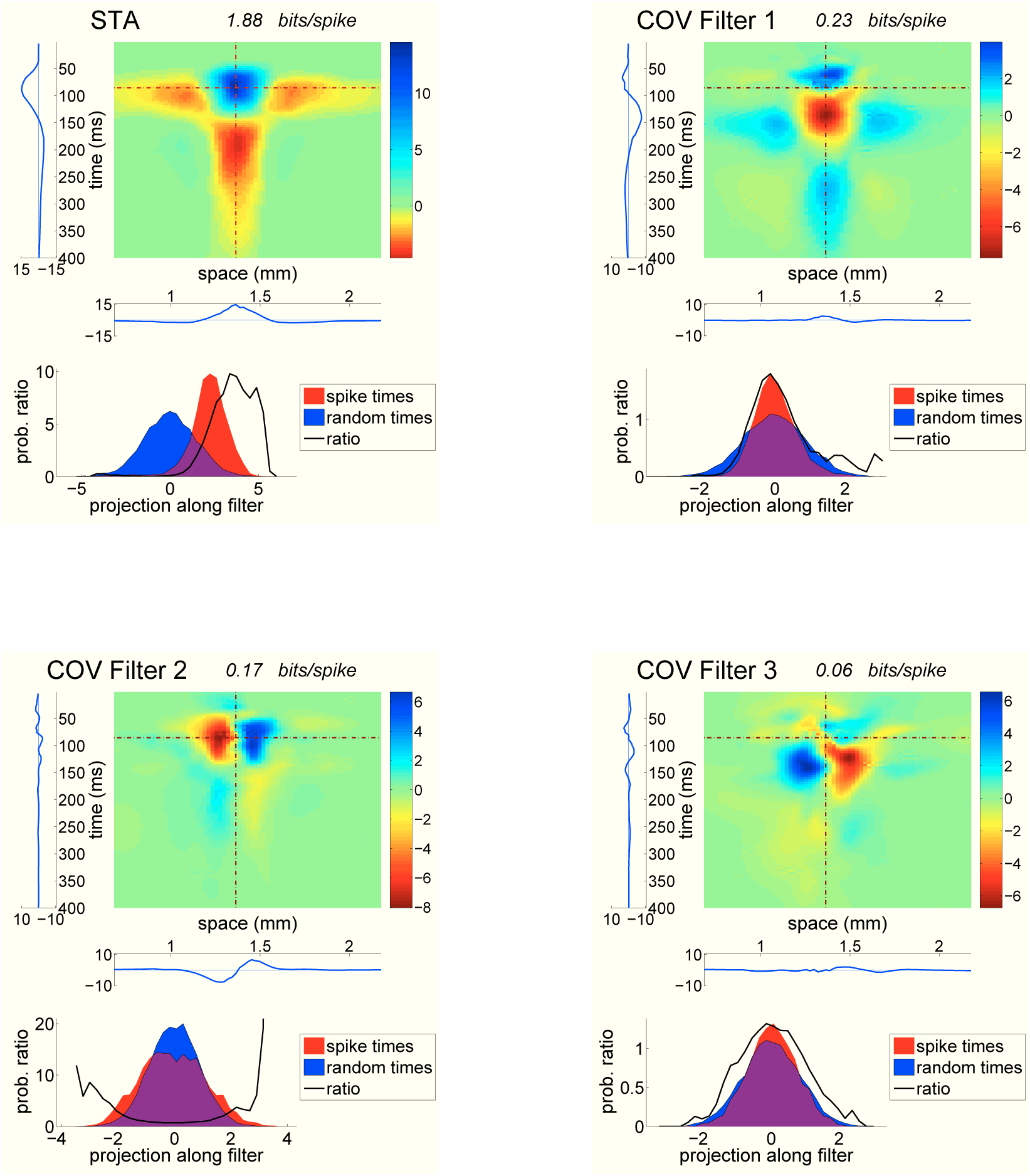
Four most informative filters for a typical brisk (biphasic-OFF) cell. Covariance filters 1, 2 and 3 are temporal edge, fast spatial edge and slow spatial edge filters respectively.

**Figure 7.**
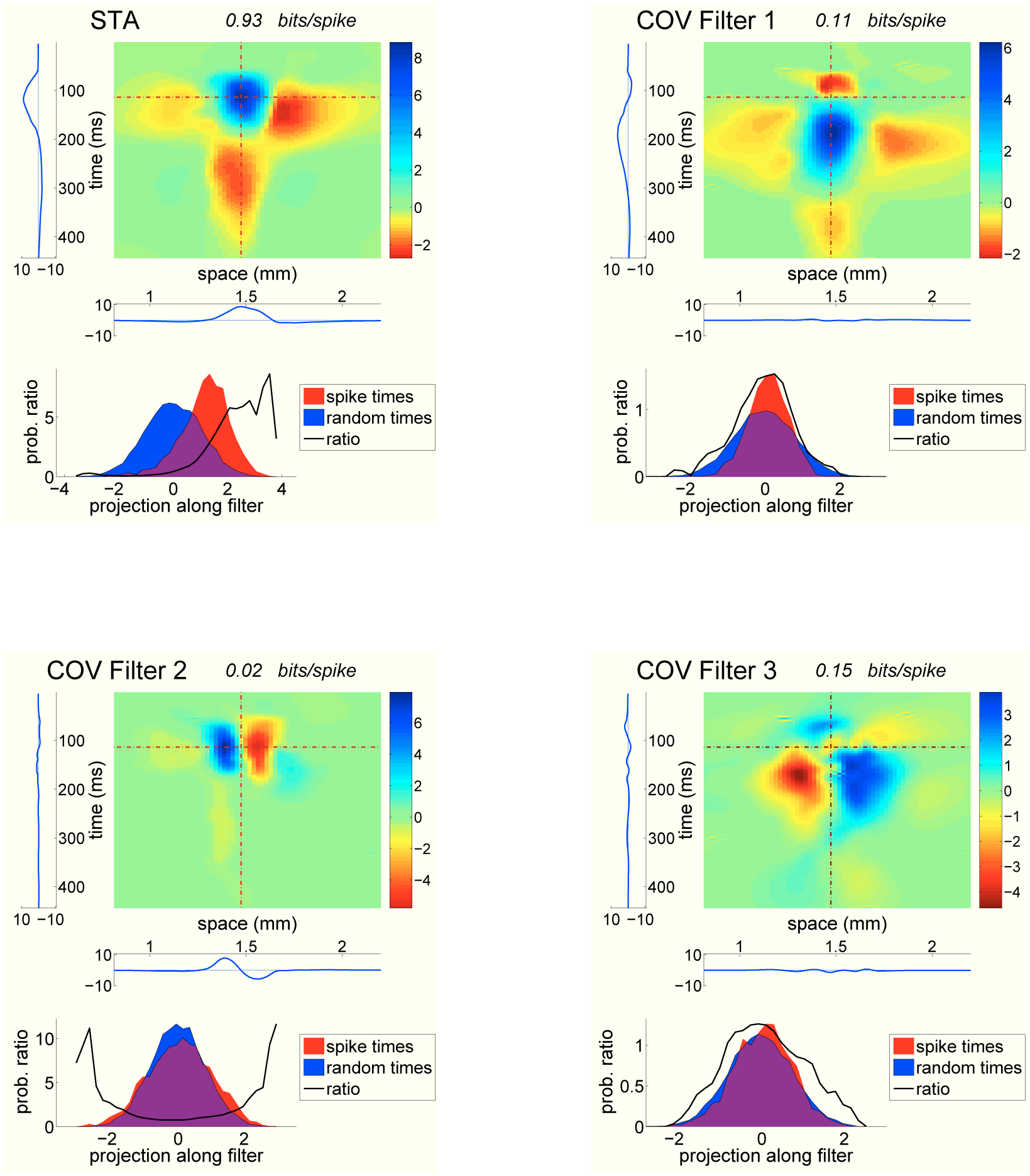
Four most informative filters for a slightly more sluggish (monophasic-OFF) cell. Covariance filters 1, 2 and 3 are temporal edge, fast spatial edge and slow spatial edge filters respectively.

Fast spatial edge filters (filter 2 in Figures 6 and 7) consist in a positive and a negative lobe, juxtaposed close to each other in space over the receptive field center, close to the time when the STA is maximal: fast spatial edges are sensitive to spatial contrasts right before a spike. Fast spatial edges are excitatory filters: the distributions of projections along fast spatial edges at spike times have roughly the same mean as the distributions of the same projections at random times, but with a significantly larger variance; stimuli with either large positive or large negative projections along the fast spatial edge elicit spikes with higher probability.

Slow spatial edge filters (filter 3 in Figures 6 and 7) consist in a positive and a negative lobe, centered spatially close to the receptive field center, appearing much farther back in time than the STA maximum or the fast spatial edge lobes. The two lobes are broader in both space and time than the fast spatial edge lobes, and they often expand spatially beyond the receptive field center. Like temporal edges, slow spatial edges are suppressive filters: the distributions of projections along slow spatial edges at spike times have a significantly smaller variance than the distribution of projections at random times; cells are more likely to fire when the stimulus projection along a slow spatial edge filter is small.

The most stereotypical spatial and temporal edge filters appeared for brisk cells. Sluggish cells gave rise to similar filters, with some variation in filter shapes. For some sluggish cells, several fast spatial edge filters appeared, with spatially shifted lobes (Fig. 8). For many sluggish cells, fast and/or slow spatial edges had more than two lobes. Despite these variations, classifying most filters as fast or slow spatial edges or temporal edges remained possible, including for the 3 on-type cells in our datasets. The covariance analysis filters which did not fall into these three categories were mostly attributable to an insufficient number of spikes for some cells.

**Figure 8.**
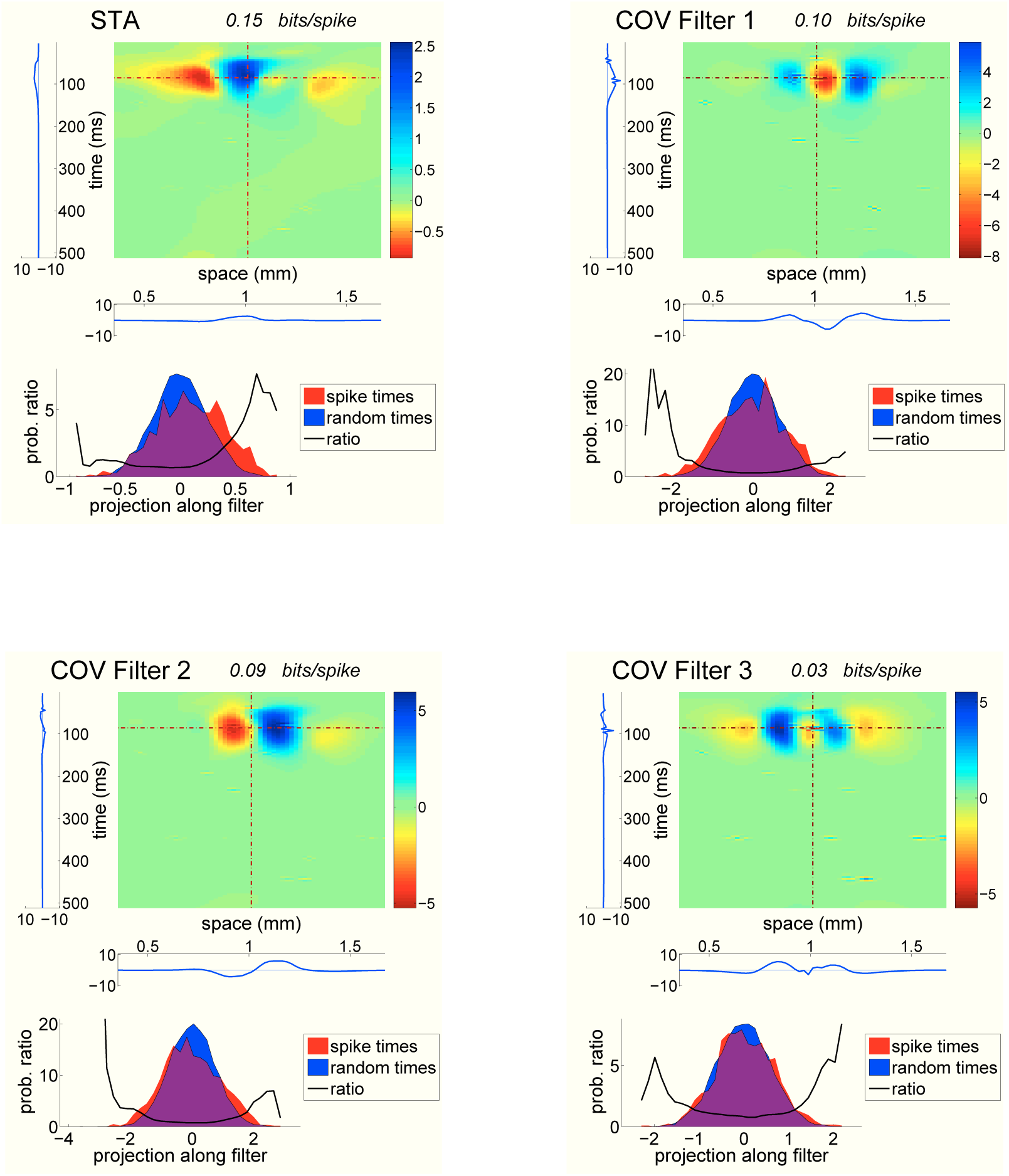
Four most informative filters for another monophasic-OFF cell. All covariance filters are fast spatial edge filters, in a broadly defined sense.

**Figure 9.**
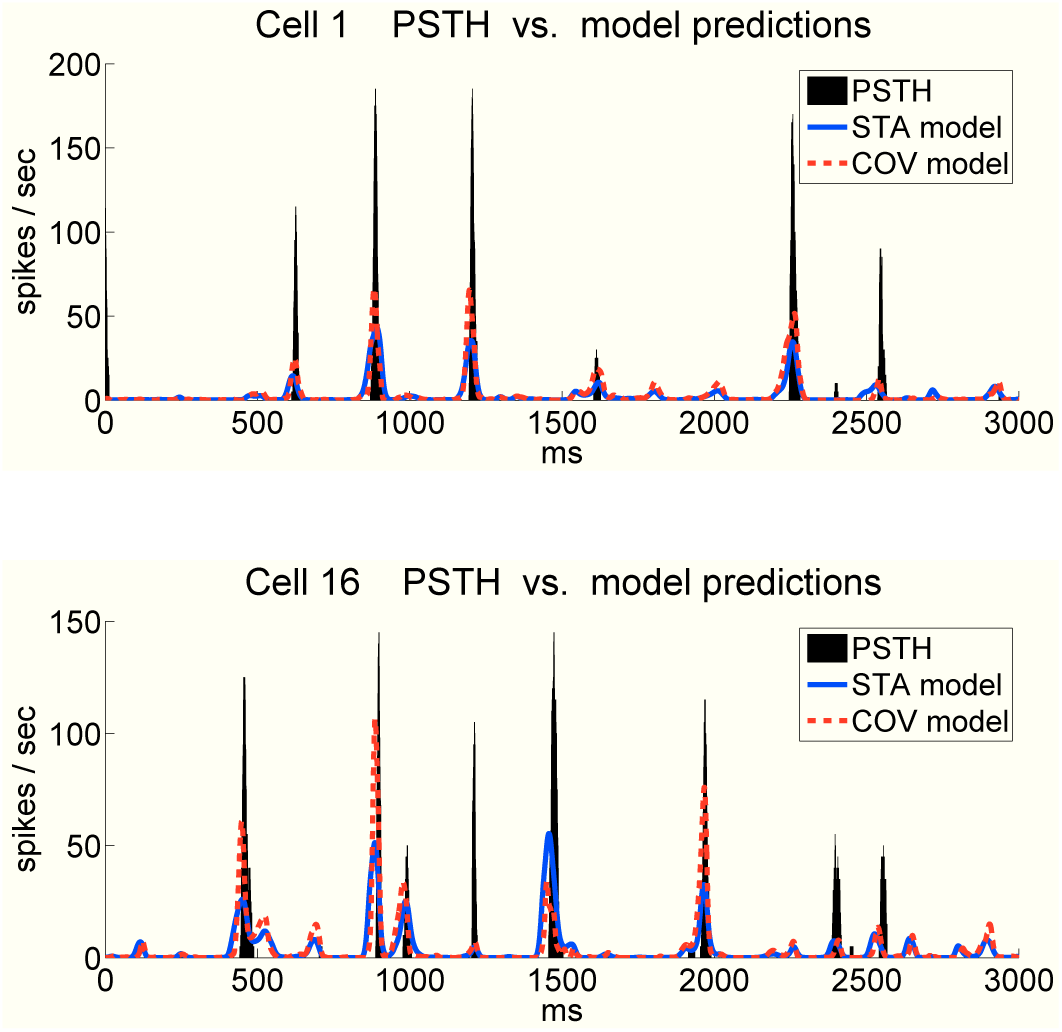
STA and COV prediction of the firing rate of two cells, overlaid against their true post stimulus time histogram (PSTH), as measured with 20 repeats, smoothed with a square window of width 5 *ms*. The COV model in both cases consisted in the STA filter augmented with 3 filters found by covariance analysis.

### Nonlinearities and information captured

We fit nonlinearities which transform our linear projections into a one-dimensional predicted instantaneous firing rate by fitting the probability densities of projections at spike times and at uniformly random times, and taking the ratio of the two. This was done on the output of each linear filter separately, as well as on several filters jointly, using kernel density estimation (see Methods).

We evaluated the information captured by the nonlinear output as well as by the linear filters directly. The amounts of information reported were estimated on a data set put aside for testing purposes only. When we compare the information captured by the nonlinear model using the best subset of the 4 most informative filters with the best direct information estimate using the same 4 filters, the amounts obtained are in rough agreement, with the nonlinear models having a slight advantage. Like other Kullback-Leibler divergence estimators, the nearest-neighbor based estimator we used has a negative finite-sample bias that becomes more prominent for higher dimensions and fewer data points: our direct multi-dimensional estimates of information captured are smaller than the true information quantities. Similarly, if the fitted nonlinearities were perfect, the amount of information captured by the output of the nonlinearity should be equal to the amount of information captured by the linear filters feeding into the nonlinearity. In practice, it is impossible to fit the nonlinearity perfectly with the amount of data available, and the information captured by the nonlinear output is in general smaller than the true information captured by the linear filters directly. Since the information estimation for the nonlinear model is simply that of a one-dimensional quantity, the information estimation itself is reliable, and the amount of information quoted for the nonlinear model is a reliable lower bound on the linear filters’ information content. The fact that this lower bound is close to the direct multi-dimensional estimates of the filters’ information suggests both that the nonlinearities are indeed well fit, and that the direct estimates are not very far off the mark. In the end, while the amounts of information cited here are the highest we could reliably ascertain with the amount of data available, they are likely to be slightly lower than the actual information content of the linear filters involved.

For comparison, we also evaluated the information captured by nonlinearities which were factorized along each dimension. Factorized nonlinearities consisted in products of nonlinearities found by kernel density estimation using the output of each linear filter separately. Interestingly, the amount of information captured by factorized models was very similar to the amount captured by the non-factorized nonlinearities. Also, the amounts of information captured by multiple filters about the stimulus were in general close to being the sum of information captured by individual filters. These two observations indicate that the contribution of each linear filter is neither clearly synergistic nor redundant: filters contribute roughly independently, and nonlinearities compose multiplicatively along features. This contrasts with the case of models including past spike histories, where the contribution of the STA and filters found by covariance analysis are clearly redundant (see).

In general, the information captured by our best models for each cell spans a broad range, from 17 to 88% of the information bound for that cell. It is notable that the cells which are captured best tend to be brisk cells, with relatively fast, spatially fine, more classical receptive fields (Figures 6, 7). For these cells, the STA and GLM filter already capture a relatively large proportion of the information bound. On the other hand, the cells which are hardest to capture with the methods presented here tend to be sluggish cells, with slow receptive fields, and large centers and surrounds.

## Discussion

We have fit models of retinal ganglion cell responses to spatially and temporally fine grained random stimuli with an emphasis on benchmarking how close our model predictions are to the true responses. The main tool which allowed us to deal with a very high dimensional stimulus space considering the relatively small amount of data available was a wavelet projection: this projection allowed us to find several informative linear filters without over-learning. For some cells, we capture more than 80% of the mutual information between cell responses and the stimulus using models which augment the STA with additional linear filters found by covariance analysis. While the supplemental information provided by each additional filter is modest, the effect of each filter is roughly independent of the others, and together their contributions add up to a significant improvement over simple STA-based models. For the most part, the additional filters can be easily classified into 3 types, with stereotyped effects on a cell’s probability of firing, offering some insight into effects that appear to be ubiquitous in retinal ganglion cell responses.

The three new types of easily recognizable filters we see in addition to the traditional STA filter let us speculate on the nature of the features being detected by cells in addition to the classical receptive field. The characteristics of the temporal and fast and slow spatial edge filters which we find for most cells are consistent with the idea that spikes mark the moment in time when a fine grained spatial contrast appears relatively still within the receptive field center: the preference for large deviations along the fast spatial edge within the receptive field center selects for a localized high frequency spatial contrast close in time to the spike ; at that same time the preference for small deviations along the temporal edge selects for little change in the overall light intensity ; the slow spatial edge selects for little spatial contrast farther in time from the spike. A stimulus which might excite a cell along all these features could be a local spatial contrast which was moving across the retina, and decelerates to a stop on the cell’s receptive field center. Another effective stimulus could be a local spatial contrast which starts out very blurred and is progressively un-blurred to reveal its contrast. Such selectivity seems likely to be desirable, since for the finer details of a scene to be detectable, the scene being projected onto a cell’s receptive field center should be properly focused and it should not be moving much faster than the ratio of the cell’s spatial to temporal integration scales. The argument that cells are locally selective for such features, as opposed to a more wide-field detection of immobility and sharpness of image shared among a population of ganglion cells, can be made because the stimulus we used does not have wide-field spatio-temporal correlations, although this does not exclude wide-field effects working as well. On the other hand, this effect is weak, and the total amount of information captured by the spatial and temporal edges do not in general exceed one half of a bit. This line of thought begs for the design of experiments which minimally excite cells along their STA feature, but instead probe the behavior of cells along the three new types of features found here.

(Ölveczky et al., 2003; Baccus et al., 2008) observe that salamander fast-OFF ganglion cells (biphasic-OFF in our setting) can detect motion in the 800*µ*m closest to the cell’s receptive field center if this motion is different from that of the background. They propose rectifying subunits, identified as being the output of bipolar cells, as the basis for this detection, both in the cell’s center and peripheral receptive field. The stimulus we used does not have long-range spatio-temporal correlations in directions where it varies, and biphasic-OFF cells being the most brisk cells are already relatively well modeled by their STAs alone, and are the cell type best captured by our models. The fast and slow spatial edges which we find for these cells are likely to be related to the rectification of subunits reported, although our analysis does not isolate individual subunits. What is more, we find that spatial edge features are informative for all the types of cells we recorded from, suggesting that most ganglion cells might be receiving rectified subunit inputs.

While they can be good predictors of some cells’ responses, STA and covariance analysis based models make for poor descriptions of many other salamander retinal ganglion cell responses. Even for the most accurate model type we tried, less than half of the information bound is captured for roughly half of the cells modeled. While this could be alleviated to some extent by having more data, as suggested by the top-right panel of Figure (10), having more data would likely not help much with the harder sluggish cells. This contrasts with the case of spatially uniform stimuli, where 2 filters found by covariance analysis have been shown to capture most of the predictable information (Fairhall et al., 2006). This suggests that many cells have many nonlinear spatial summation properties on a fine scale which confound our mostly linear analysis methods. This is corroborated by the fact that for many cells, several spatial edge filters can be found, allowing us to speculate about what our present models are missing: at play are likely many more than the 2-3 spatial edge filters distinguishable with the limited amount of data available. This is all the more intriguing that many of the cells with several spatial edge filters are the sluggish cells which are being most poorly modeled.

**Figure 10.**
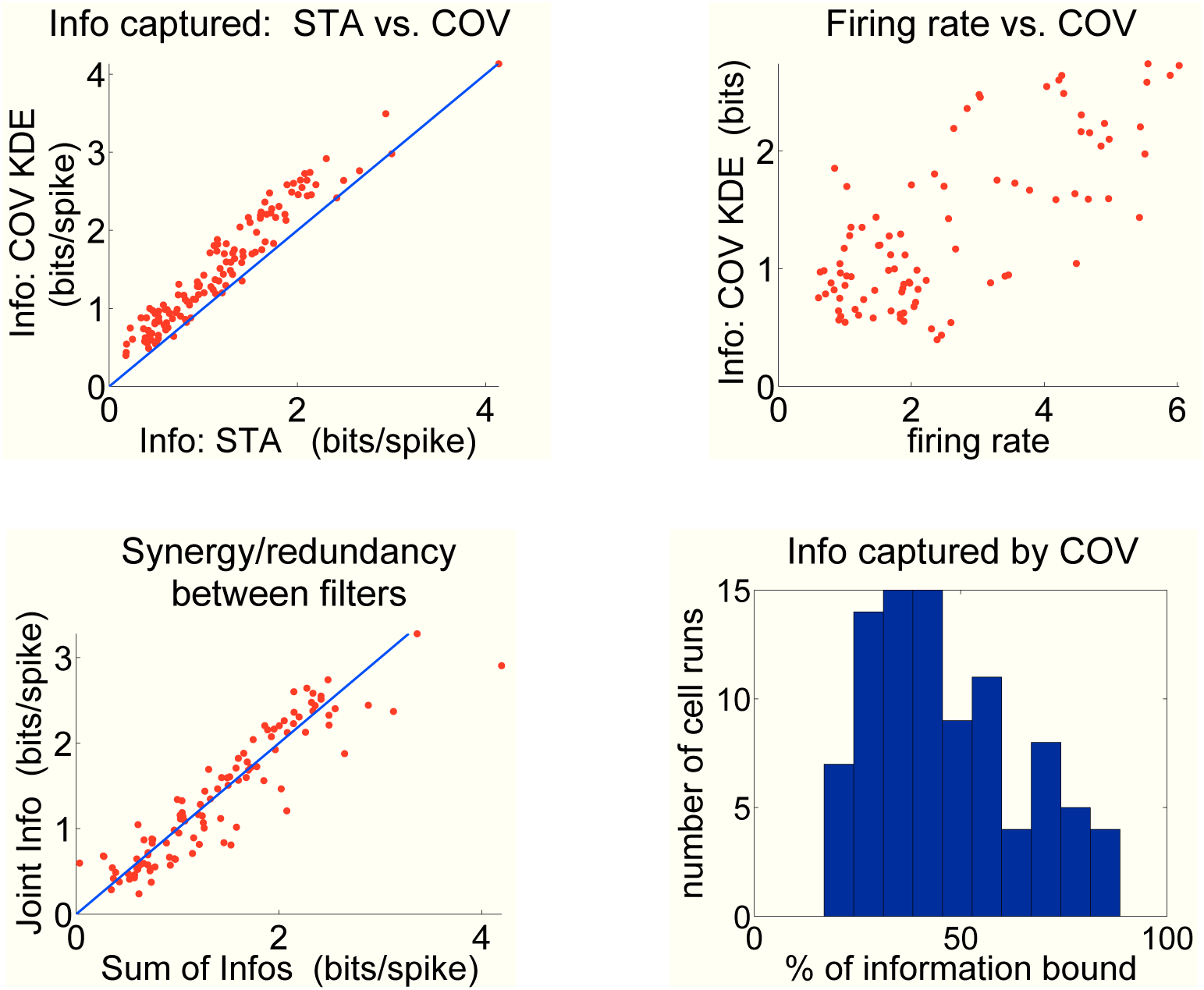
Percentage of the information bound ℬ captured by the STA *(top left)* and firing rate *(top right)* plotted against the information captured by the STA filter supplemented with at most 3 filters found by covariance analysis, with a nonlinearity estimated by kernel density estimation (‘COV KDE’). *Bottom left:* comparison of the sum of informations captured individually by 4 filters with the information captured by the 4 filters jointly. The information that filters carry about the stimulus is neither clearly redundant nor synergistic. *Bottom right:* histogram of the percentage of the information bound B captured by the COV KDE model.

**Figure 11.**
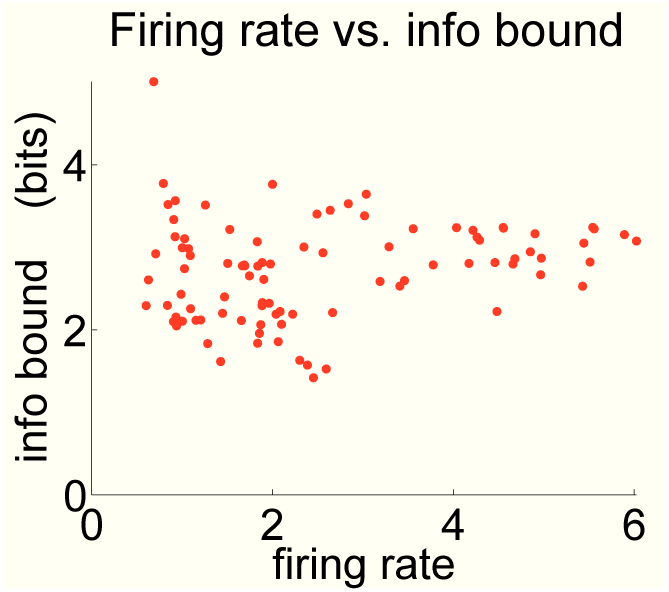
True Brenner information as evaluated from 20 repeats of 30 seconds of random stimulus, using the nearest-neighbor estimator.

**Figure 12.**
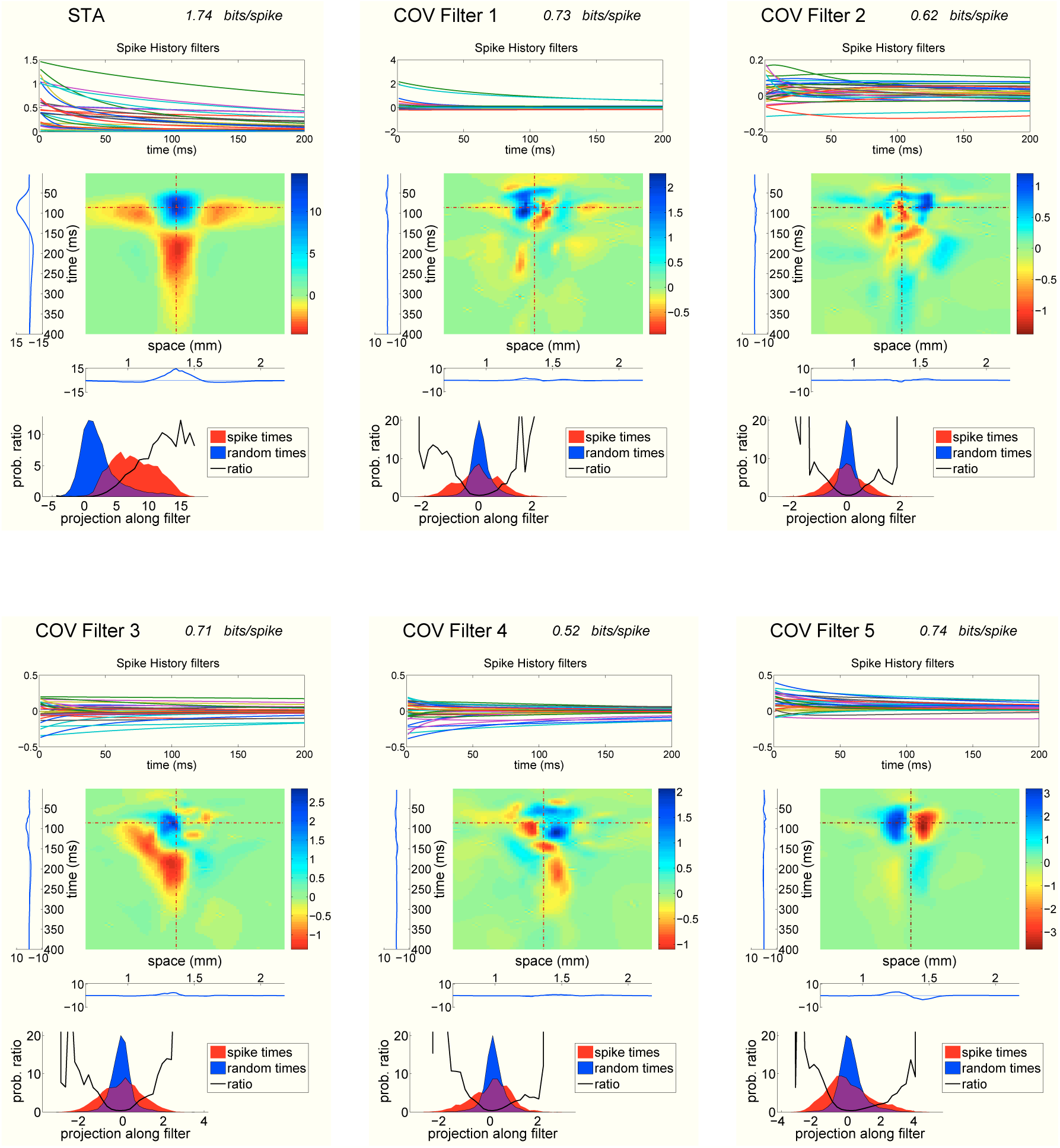
Six most informative filters for a brisk (biphasic-OFF) cell. Covariance filters 1-4 seem to be dominated by their spike history components, as their stimulus history components seem garbled.

## Supplementary Material

### Adding spike history dependence

We looked at the statistical structure present in spike trains that is not explainable through stimulus dependencies by including spike histories alongside with the past stimulus in our models. Spike histories were calculated using the time since the last spike for all the cells in the population (*n* = 23 and 37 respectively for each experiment). The spike history dependence for both the GLM (see the next section) and covariance analysis filters was parameterized by the family of exponential functions exp(− *s*_*c*_(*t*)*/τ*_*j*_), for *τ*_*j*_ ∈ {1, 2, 3, 5, 8, 13, 21, 34, 55, 89} *ms*, where *s*_*c*_(*t*) is the time since the last spike of cell *c* at time *t*. Our covariance analysis methods were applied unchanged to the set of pasts consisting of the concatenation of the wavelet coefficients characterizing past stimuli and the vector representing the times since the last spike of each cell by applying the exponential functions exp(−*s*_*c*_(*t*)*/τ*) for each *τ*_*j*_ and each cell *c*.

Two differences with the stimulus only case are to be noted in the results obtained. Firstly, adding information about past spikes invalidates the upper information bound ℬ on the amount of information captured by the model. Secondly, the addition of spike histories is likely to make covariance analysis less effective. This is because in the case of models with past stimuli only, the distributions of projections along the filters found were approximately gaussian and mostly unimodal, so that picking out filters with large differences in variance was an effective way of finding informative dimensions. For models with spike history dependence on the other hand, projections along filters tend to be multimodal and far from gaussian, and covariance analysis is less apt at picking out informative dimensions. In particular, adding spike histories should in theory always increase the total information captured, while this is not always the case with our methods (Figure 13, middle panel). This suggests that our covariance methods are not ideal for including spike train histories. Nonetheless, we briefly describe in the next paragraph the results obtained.

**Figure 13.**
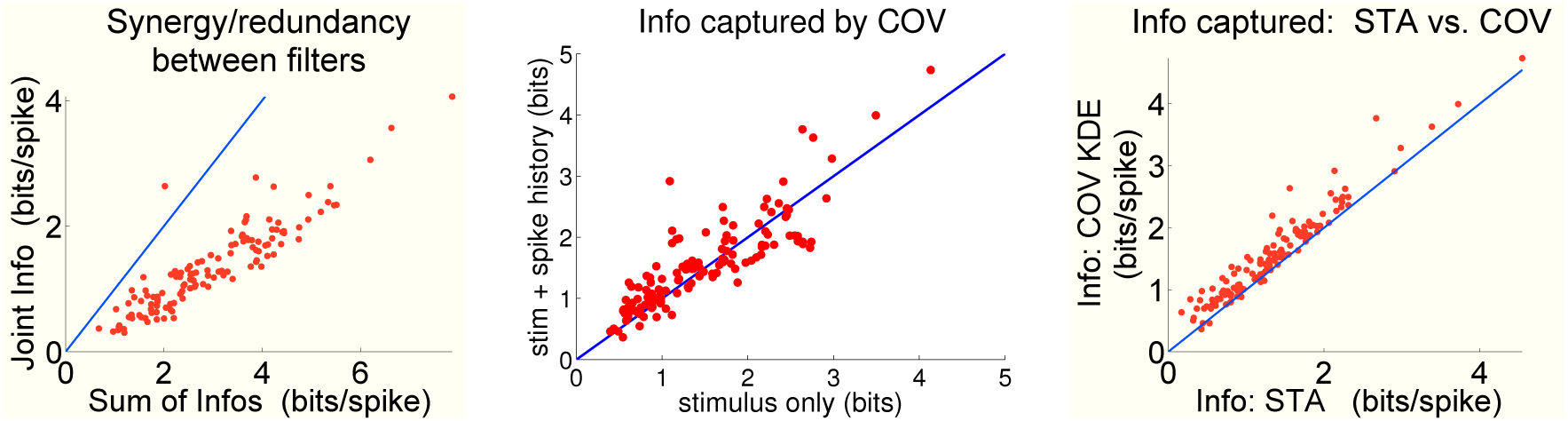
Information captured by covariance models including spike histories. *Left:* comparison of the sum of informations captured individually by 4 filters with the information captured by the 4 filters jointly. The information that filters carry about the stimulus is clearly redundant. *Middle:* comparison of information captured by the best covariance model with and without spike history dependence. *Right:* Information captured by the STA compared with the information captured by the best covariance model.

**Figure 14.**
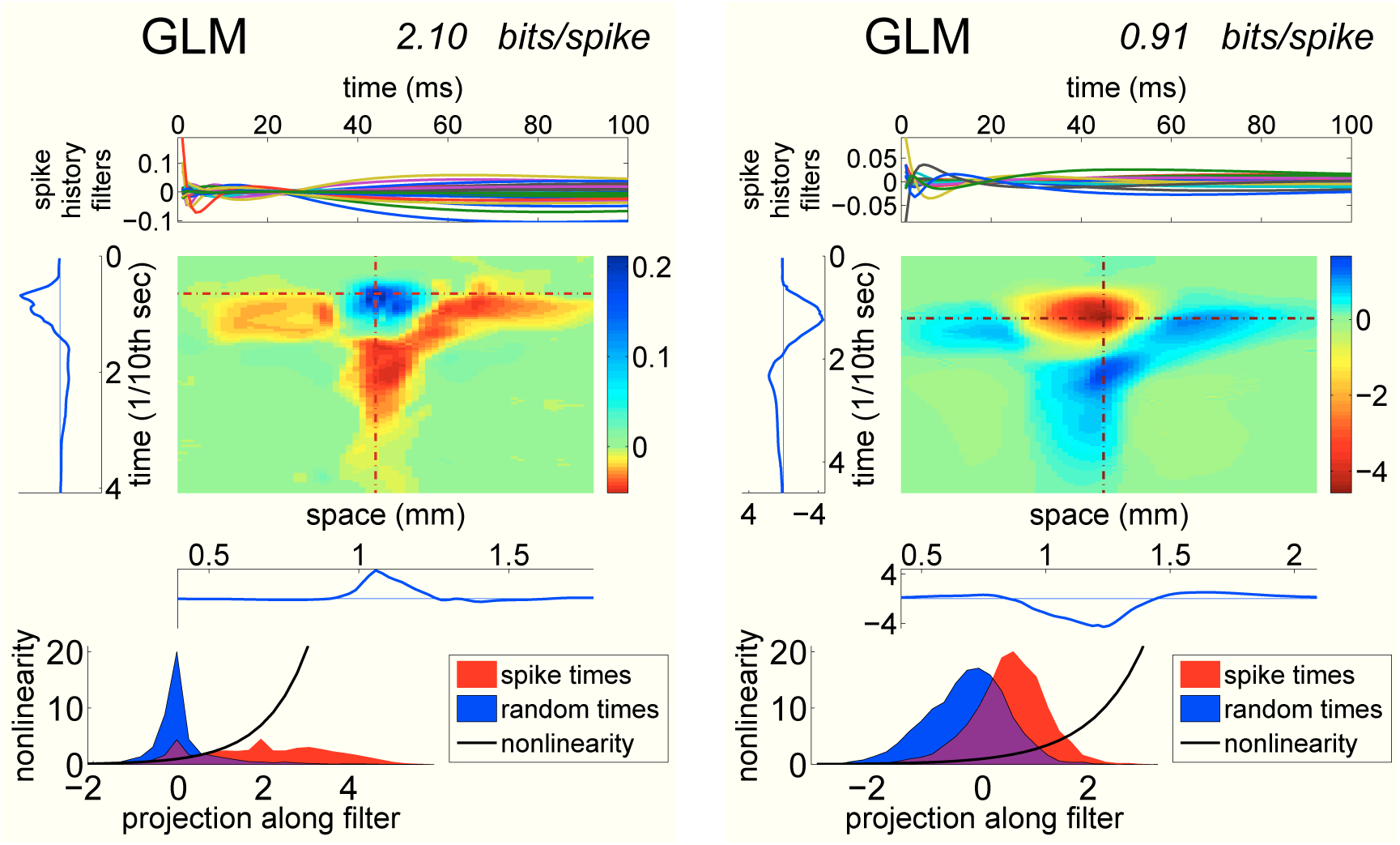
GLM filters with spike history dependence for a brisk cell (biphasic-OFF, left) and a fast-ON cell (right).

**Figure 15.**
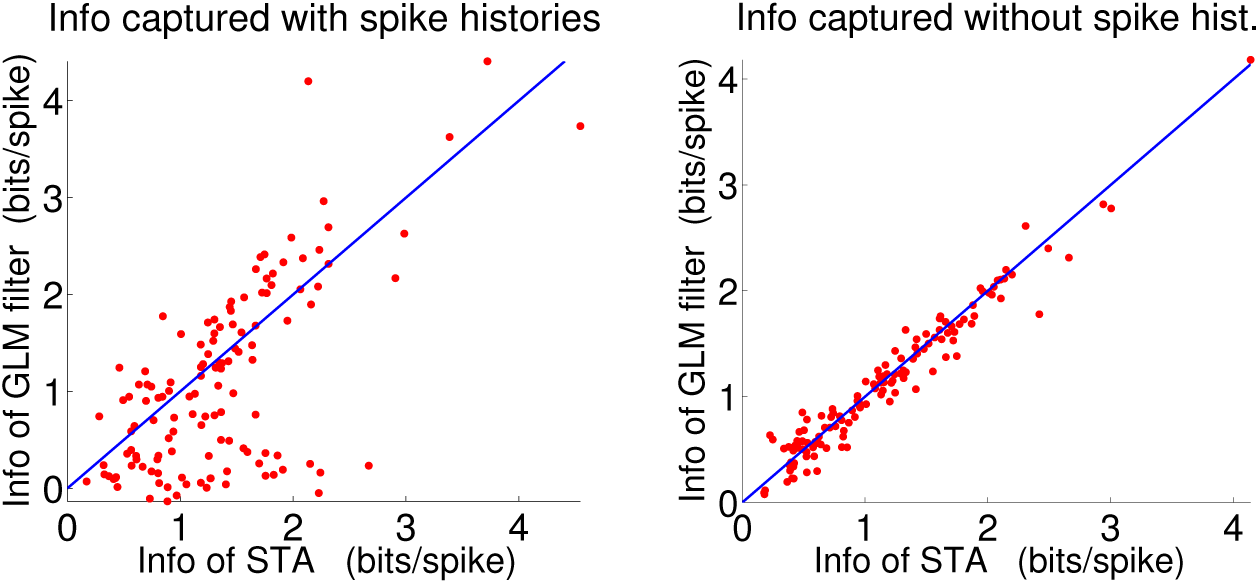
Information captured by the STA plotted against the information captured by the GLM filter.

In general, adding spike histories typically produces 6 or more covariance filters with large information content, as opposed to the 2 or 3 found without spike histories. Many of the resulting covariance filters are dominated by the contribution of the spike histories, with stimulus history filters looking mostly like noise, while for other filters, we obtain smooth stimulus history components that often fall into the categories found without stimulus histories: STA, temporal and spatial edges. A large fraction of covariance filters have distributions of projections which have larger variance at spike times than at random times, as with the fast spatial edge filter. The spike history filters are often large for many cells, and more often than not, the cell’s own spiking history filter is no larger than that of other cells in the population. Unlike models without spiking histories, the information that linear filters including spike histories carry about the stimulus are clearly redundant (Figure 13, left panel).

### Generalized Linear Models

Generalized Linear Models (GLM) have been very successful at modeling macaque parasol ganglion cell responses to random stimuli (Pillow et al., 2008). GLMs are linear-nonlinear models with a single linear filter and an exponential nonlinearity, whose output is interpreted as a spiking intensity, as in (Siebert, 1965; Bialek & Zee, 1990). Like our other models, the GLM linear filter is applied to the wavelet coefficients representing past stimuli and optionally spike train pasts. Its parameters are obtained for each cell by maximizing the log-likelihood *L* of some spike train data under the model using standard quasi-newton optimization algorithms, where *L* = ∑log *λ* (*t*_*sp*_) − ∫*λ* (*t*) *dt*. The conditional intensity of a given cell’s spiking activity was assumed to be of the form *λ*(*t*) = exp(**k**.**x**+ ∑_*c*_ *l*_*i*_(*s*_*c*_(*t*))+*µ*), where **x** are the wavelet coefficients at time *t, s*_*c*_(*t*) is the time interval between the last spike of cell *c* and *t*, and *µ* is a baseline log-firing rate. *l*_*c*_(*t*) is the linear filter applied to the time *s*_*c*_(*t*) since the last spike of cell *c*, and it was expressed as a weighted sum of exponential functions exp(−*t/τ*_*j*_) as in the previous section. We fit two types of GLM models to our data: models depending on the stimulus wavelet coefficients only (*l*_*i*_ = 0) and models depending on stimulus wavelet coefficients as well as past spikes from all cells in the recording. Adding a regularization penalty to the log-likelihood was essential to avoiding over-fitting. We used an *L*_2_ penalty 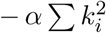 on the wavelet parameters **k** and the penalty 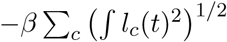 on the spike history filters *l*_*c*_. The regularization parameters *α* and *β* were chosen to maximize the model likelihood over a validation data set.

The GLM model filters we obtain resemble each cell’s STA in stimulus space, whether or not past spikes were included. For models without spike histories, the amount of information captured by the GLM filter was close to the information captured by the STA. This is most likely due to the fact that although regularizing the GLM filter was essential to avoid over-fitting, the resulting smoothing of filters means less information is being captured. For models depending on the past stimulus only, the proportion of Brenner information captured varies between 5 and 70% depending on cells. Many cells are thus being modeled quite poorly, as with the STA.

The information captured by GLM models with spiking histories were less closely related to the information captured by STA models, and could be more or less informative than the STA depending on cells: for some cells, the GLM filter was a clear improvement over the STA, while for others the STA was clearly better. In any case, the GLM spike history filters capture more of the fast timescale structure close to the spike, whereas the STA and covariance filters tend to have more slowly varying spike history filters.

### Mutual Information calculations

Let us see how one can express the mutual information between spikes {*t*_*i*_} and a arbitrary quantity *X*(*t*). In practice, the kinds of *X*(*t*) we consider are functions of the stimulus before time *t* that carry information about whether or not there was a spike at time *t*; we could also take *X*(*t*) to be the whole past stimulus before *t*. Let us consider the infinitesimal probability that a spike occurred in the interval [*t, t* + Δ*t*[. By definition, the mutual information **MI**(*spike* ∈ [*t, t* + Δ*t* [; *X*(*t*)) is written as the following integral over the domain of *X* and whether or not a spike occurred in [*t, t* + Δ*t*[:

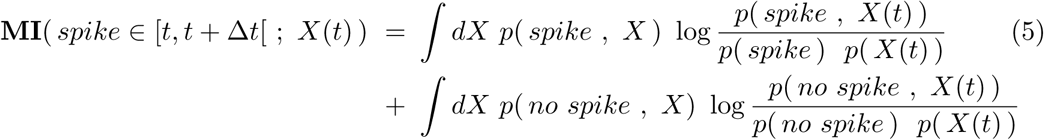

We can apply Bayes rule in two different ways to obtain two different points of view on how to calculate relevant mutual informations empirically. In both cases, we will see that the *no spike* term in (5) is zero.

First, let us condition on *X*. In the limit where Δ*t* becomes small, we can write the first order approximation *P* (*spike* |*X*) ≈ *r*(*X*)Δ*t*, where *r*(*X*) is the instantaneous spiking rate given *X*. Similarly, writing < *r* > to denote the average of *r*(*X*) over *p*(*X*), we have *P* (*spike*) ≈ < *r* > Δ*t*. The mutual information (5) then becomes:

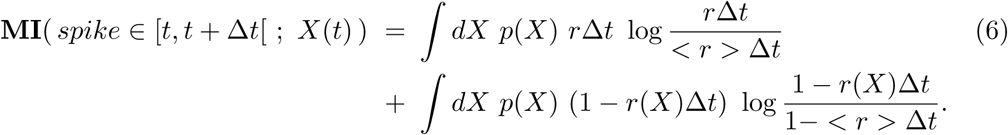

As it turns out, the second term vanishes as Δ*t*^2^: to first order, the second integral is equal to ∫*dX p*(*X*) (< *r* > *r*(*X*))Δ*t*, which by definition of < *r* > is zero. The mutual information is thus dominated by the first term, which vanishes as Δ*t*. Assuming the joint statistics of *X*(*t*) and the spike train to be stationary, it makes sense to divide this vanishing mutual information by < *r* > Δ*t* to obtain a mutual information rate ℐ(*spike, X*(*t*)) in *bits per spike*, and to replace averages over *X* and the spike train by averaging over time:

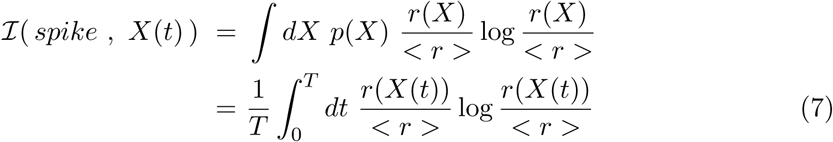

This last expression (7) can be used for an important practical calculation. If *X*(*t*) is taken to be the whole past stimulus at time *t*, we can estimate *r*(*X*(*t*)) empirically for a range of values of *X* by showing a certain snippet of stimulus many times: *r*(*X*(*t*)) is then the average firing rate at time *t* within the snippet, as estimated from all the repeats. This allows us to calculate ℐ(*spike, past stimulus*), which is an upper bound on the mutual information between spiking and any function of the past stimulus. ℐ(*spike, past stimulus*) is a measure of how noisy or unreliable a cell’s responses are.

If *X*(*t*) is some low-dimensional quantity such as a small set of filters or the output of a model designed to predict spikes, we could use this same expression (7) to estimate the mutual information between spikes and the predictor *X*. This approach has the disadvantage that it would require estimating *r*(*X*), presumably from repeats. As it turns out, we can do better by doing the other Bayes inversion of (5) where we condition on the presence or absence of a spike.

Treating the spike train mathematically as a point process, spikes have no width in time, and conditioning on the absence of a spike is equivalent to not conditioning at all. In particular, *p*(*X* |*no spike*) = *p*(*X*), which immediately implies that the second term in (5) is zero. Once again we obtain the mutual information rate ℐ in *bits per spike* by dividing **MI** by < *r* > Δ*t* = *p*(*spike*):

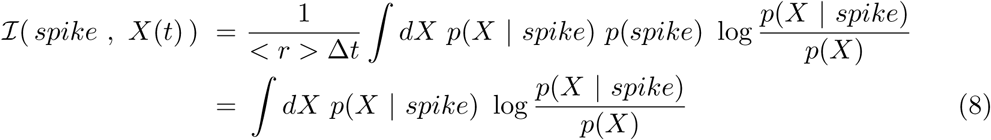

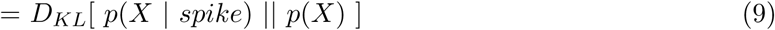

This exposes the sought mutual information rate as the Kullback-Leibler divergence between the distribution of *X* at spike times relative to its distribution over time. This calculation makes it clear that to look for functions *X* of the past stimulus which informative about a cell’s spikes, we should compare the ensemble of past stimuli before spike times with the distribution of past stimuli over time, and see how they differ.

